# Terminitor: Cleavage Site Prediction Using Deep Learning Models

**DOI:** 10.1101/710699

**Authors:** Chen Yang, Chenkai Li, Ka Ming Nip, René L Warren, Inanc Birol

## Abstract

As a widespread RNA processing machinery, alternative polyadenylation plays a crucial role in gene regulation. To help decipher its underlying mechanism and understand its impact, it is desirable to comprehensively profile 3’-untranslated region cleavage and associated polyadenylation sites. State-of-the-art polyadenylation site detection tools are known to be influenced by library preparation artefacts or manually selected features. Moreover, recently published machine learning methods have only been tested on pre-constructed datasets, thus lacking validation on experimental data. Here we present Terminitor, the first deep neural network-based profiling pipeline to make predictions from RNA-seq data. We show how Terminitor outperforms competing tools in sensitivity and precision on experimental transcriptome sequencing data, and demonstrate its use with data from short- and long-read sequencing technologies. For species without a good reference transcriptome annotation, Terminitor is still able to pass on the information learnt from a related species and make reasonable predictions. We used Terminitor to showcase how single nucleotide variations can create or destroy polyadenylated cleavage sites in human RNA-seq samples.

**Author Summary:** 3’ cleavage and polyadenylation of pre-mRNA is part of RNA maturation process. One gene can be cleaved at different positions at its 3’ end, namely alternatively polyadenylation, thus identifying the correct polyadenylated cleavage site (poly(A) CS) is essential to unveil its role in gene regulation under different physiological and pathological conditions. The current poly(A) CS prediction tools are either heavily influenced by RNA-Seq library preparation artefacts or have only been designed and tested on *ad hoc* datasets, lacking association with real world applications. In this study, we present a deep learning model, Terminitor, that predicts the probability of a nucleotide sequence containing a poly(A) CS, and validated its performance on human and mouse data. Along with the model, we propose a poly(A) CS profiling pipeline for RNA-seq data. We benchmarked our pipeline against competing tools and achieved higher sensitivity and precision in experimental data. The usage of Terminitor is not limited to genome and transcriptome annotation and we expect it to facilitate the identification of novel isoforms, improve the accuracy of transcript quantification and differential expression analysis, and contribute to the repertoire of reference transcriptome annotation.

## Introduction

Since polyadenylated (poly(A)) sequences were first discovered on the 3’ ends of eukaryotic mRNAs 40 years ago [1], numerous studies have contributed to the understanding of the mechanisms, evolution, regulation, and impacts of polyadenylation [2–5]. As an essential step in mRNA maturation, RNA polyadenylation involves two phases: 3’ end cleavage of nascent RNA and addition of a poly(A) tail. This process is facilitated by several *cis*-acting RNA elements, represented by the poly(A) signal (PAS), conserved hexamer motifs with AATAAA being the most strongest, located 10-30 nt upstream of the cleavage site (CS) [6]. Because most nascent RNA in eukaryotes have more than one possible 3’ end CS, mRNAs with different 3’ ends are formed following these two coupled processes [7]. This phenomenon is also known as alternative polyadenylation (APA), which results in transcripts with different 3’ untranslated regions (UTR). In recent years, advances in DNA/RNA sequencing technology allowed detailed study of APA modulation under different physiological and pathological conditions, and shed light on its implications in some diseases, especially cancer [5,8–10].

### The importance of profiling 3’ UTR cleavage

APA, together with alternative splicing, has considerable impact on the modulation of gene expression and contributes to transcriptome complexity. As of date, the catalogue of profiled poly(A) CS, especially cancer-specific ones, is far from complete [11]. Ensembl annotation version GRCh38.94 records 63,620 poly(A) CS for 19,907 protein coding genes in human, and among them, 73.13% genes can produce two or more APA isoforms (S1 Fig) [12]. The most recent poly(A) database, PolyA_DB3, compiled 24 human samples and cataloged 108,042 poly(A) CS for 20,998 genes [13]. The substantial improvement in 3’ end annotation compared to Ensembl suggests the sample-specific usage of poly(A) CS, highlighting the incompleteness of the current annotation. Moreover, usage of these poly(A) CS is dynamically regulated under different developmental stages and pathological changes, as elucidated in previous studies [5,8,14,15]. Profiling poly(A) CS with respect to biological conditions serves as the first step towards deciphering the underlying mechanism of APA-mediated gene regulation. Hence, it is desirable to incorporate a fast and robust poly(A) CS characterization tool into standard transcriptome analysis pipelines.

### Current methods and their limitations

In the past decades, continuous efforts have been made to annotate 3’ ends in the genome and predict poly(A) CS in the transcriptome, using methods designed for two major sequencing protocols: 3’ end targeted sequencing and poly(A)-selected RNA-seq. The former is a specialized high throughput sequencing technique that is developed to overcome the low sequence complexity at the 3’ end of transcripts. These methods, including TAIL-Seq, PolyA-Seq, PAS-seq, and PAT-seq [16–19], only interrogate the 3’ ends of transcripts by primers, followed by sequencing and bespoke analysis pipelines. Compared to methods that use RNA-seq, a critical advantage of these protocols is their high sensitivity in detecting poly(A) CS from lowly expressed transcripts. With the help of 3’ end sequencing technologies, several poly(A) CS databases have been established [13,20–22]. Though powerful, they require specialized library preparation, which can be costly and laborious, and their utility is restricted to poly(A) detection. As a result, they have not yet been widely adopted in genetic research.

In contrast, RNA-seq is the established sequencing method of choice for a wide range of transcriptomics projects. It can also be used for poly(A) CS profiling on a per-sample basis, as well as for systematic APA analysis of large cohorts of individuals, including retrospective studies. Poly(A) CS prediction and annotation tools designed for RNA-seq have demonstrated that with suitably redundant coverage (read coverage ≥ 2), RNA-seq data type contains sufficient information for the identification of poly(A) tails, and thus permits the discovery of novel poly(A) CS [23,24]. Generally speaking, these tools can be classified into three approaches: read-evidence based, expression level based, and machine learning based.

Read-evidence based approaches, such as KLEAT [23] and ContextMap2 [24], search for untemplated adenosines (As) in aligned sequences as evidence to determine the location of poly(A) CS. Although these tools are able to discover novel poly(A) CS with high resolution, they may lose sensitivity in low coverage sequencing libraries and when transcript expression is low.

For approaches based on expression levels, poly(A) CS prediction is a by-product of APA analysis. Tools adopting this approach, which include DaPars, APAtrap, and QAPA [25–27], all rely on differential expression in the 3’ UTRs of matched samples. However, due to the low sequence complexity of 3’ UTRs and the resulting sequencing data artefacts, read coverage in these regions may not reflect true expression levels, impacting differential expression analysis and associated statistical tests. Consequently, these methods may fail to detect real but lowly represented APA events. Moreover, they only report poly(A) CS involved in significant APA events through comparative analyses, and thus these tools are not capable of profiling poly(A) CS in given samples.

Another approach to address the prediction problem is through the use of deep learning methods, and several such models have been proposed recently, such as DeeReCT-PolyA [28], DeepGSR [29], and DeepPASTA [30]. For these tools, hidden features are learnt and modeled directly from input nucleotide sequences. Using this approach, DeeReCT-PolyA and DeepGSR have been reported to show around 90% accuracy on test sequences [28,29]. Combining both sequence and RNA secondary structure information, DeepPASTA, was reported to have an area under the curve score over 93% in predicting poly(A) CS. However, these tools were only bench-marked on pre-constructed datasets composed of positive and negative genomic sequences, and none have been applied to real experimental transcriptome data for prediction. Another limitation of DeeReCT-PolyA and DeepGSR is that they were only trained to distinguish whether a PAS is real. However, it is known that a small fraction (yet a substantial number) of functional human poly(A) CS does not require a PAS [32,33]. At present, these approaches overlook poly(A) CS with noncanonical *cis*-acting elements, which would likely negatively impact their performance on experimental data.

### Terminitor

Despite the availability of various tools, comprehensively and efficiently profiling poly(A) CS in RNA-seq samples remains challenging. In this study, we present Terminitor, a deep neural network (NN) model for fast and accurate poly(A) CS recognition, independent of the existence of a PAS. Terminitor takes fixed length sequences as input and performs a three-label classification to determine whether the sequence contains a poly(A) CS, a non-polyadenylated CS, or non-CS. Coupled with Terminitor, we also propose a profiling pipeline that generates prediction of poly(A) CS using raw RNA-seq data as input. The performance of this pipeline was cross-validated on two datasets of sequences (describing human and mouse transcriptomes) to select the optimal input sequence length and model hyperparameters. We have benchmarked our pipeline against competing tools on experimental paired-end RNA-seq libraries generated from the Illumina (San Diego, CA) TruSeq paired-end and Pacific Biosciences (Menlo Park, CA) Single Molecule, Real-Time (SMRT) Sequencing platforms.

## Results

### Performance of Terminitor using different sequence lengths

Terminitor is trained on experimentally validated annotated nucleotide sequences containing three classes of sites: (1) poly(A) CS, (2) non-poly(A) CS, and (3) non-CS. Using cross-validated optimal hyperparameters, it computes the probability of a test sequence belonging to one of the three classes.

To train, validate, and test the model, we collated two reference datasets describing human and mouse transcriptomes, based on poly(A) CS databases and Ensembl annotations. These datasets are composed of labeled sequences of uniform length with three parts: upstream sequence, a site from one of the three classes, and downstream sequence. Upstream sequences are from corresponding transcripts for the first two classes and from genomic regions for the third class, while downstream sequences are genomic, and the transition from upstream sequence to downstream sequence is marked as the cleavage site or lack thereof (S2 Fig). Since there is no consensus from previous work on how the input sequence length influences prediction accuracy, we built models with different lengths to evaluate its effect on performance. Eight upstream lengths (200, 160, 120, 100, 80, 60, 40 and 20 nt) and five downstream lengths (200, 100, 40, 20 and 0 nt) were chosen to create a total of 40 models, which were trained to explore the impact of sequence length on poly(A) CS prediction (Fig 1). In general, all models show clear separation between poly(A) CS and the other two labels (S3 Fig). The accuracy of all 40 models is above 87%, including the one trained with only 20 nt long upstream sequence. As expected, models trained with sequences containing both upstream and downstream sequences perform better than the ones trained with upstream sequences alone (0 nt downstream sequence length). However, given the same upstream sequences, models trained with downstream sequences longer than 20 nt perform comparably to each other. The 28 models trained with upstream sequence lengths in {40, 60, 80, 100, 120, 160, 200} and downstream sequence lengths in {20, 40, 100, 200} have an average accuracy of 94.50% ± 0.0015. More interestingly, longer downstream sequences do not necessarily lead to better performance. Models trained with 200 nt downstream sequences have slightly lower accuracy than the ones trained with shorter lengths, suggesting that some polyadenylation recognition patterns maybe obscured by sequence motifs further downstream.

**Fig 1.**
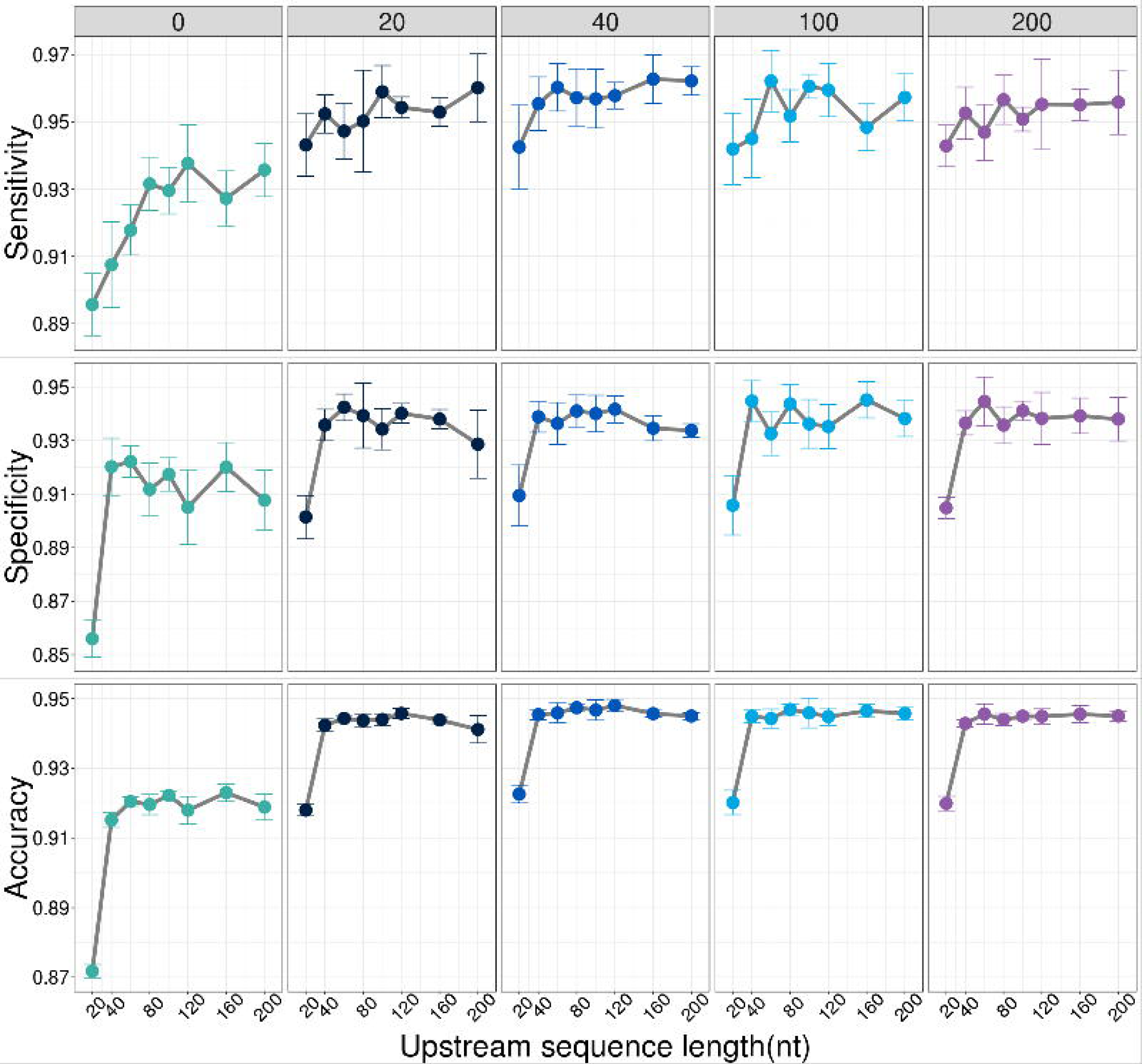
Model performance on human reference dataset. Each facet represents one downstream sequence length, and the x-axis of each facet plot shows the up-stream sequence length. Each data point is the mean of 5-fold cross validation and the error bar is the standard deviation.

On the other hand, given the same downstream sequence, the performance of models plateaus when the upstream sequence is 40 nt or longer. Similarly, longer upstream sequences do not guarantee better performance either, especially when the upstream length is much longer than the downstream length. For example, the model with 120 nt upstream and 20 nt downstream sequences is nearly 0.5% more accurate than the one with 200 nt upstream and 20 nt downstream sequences. This decrease is mainly due to a drop in specificity, which means the model has more false positive predictions with longer upstream sequence.

### Tool comparison on experimental data

In order to demonstrate the application of Terminitor on poly(A) CS profiling, we designed a prediction pipeline and applied it on two human RNA-seq samples, Human Brain Reference RNA (HBR) and Universal Human Reference RNA (UHR), to predict poly(A) CS from reconstructed transcripts. As done previously, we trained Terminitor with 40 length combinations, applied all 40 models on UHR sample, and computed the Pareto frontier based on sensitivity and precision. The sensitivity of Terminitor is positively correlated with upstream sequence length, while there is no clear trend between the downstream sequence length and the performance metrics (S4 Fig). For demonstration purposes, we chose ± 100 nt model for performance benchmarking and comparison.

Profiling poly(A) CS for species without a good reference annotation can be challenging. Thus it is interesting to test whether the information learnt from one species is transferable to another. In addition to the model built on the above-mentioned human reference dataset, we also generated a model that was separately trained on the mouse dataset. We applied these two models on the UHR and HBR samples and compared the performance with KLEAT, DeepPASTA, DeeReCT-PolyA and DeepGSR (Fig 2A, S5A Fig). Poly(A) CS predicted by these methods were compared against Ensembl annotation to calculate the sensitivity and precision. Note that the denominator in sensitivity calculations represents all Ensembl annotations, yet for a given sample not all transcripts would be expressed. Without a *priori* information on gene and isoform expression, this is the most appropriate solution at it penalizes all methods the same way, preserving their relative performance for fair comparisons.

**Fig 2.**
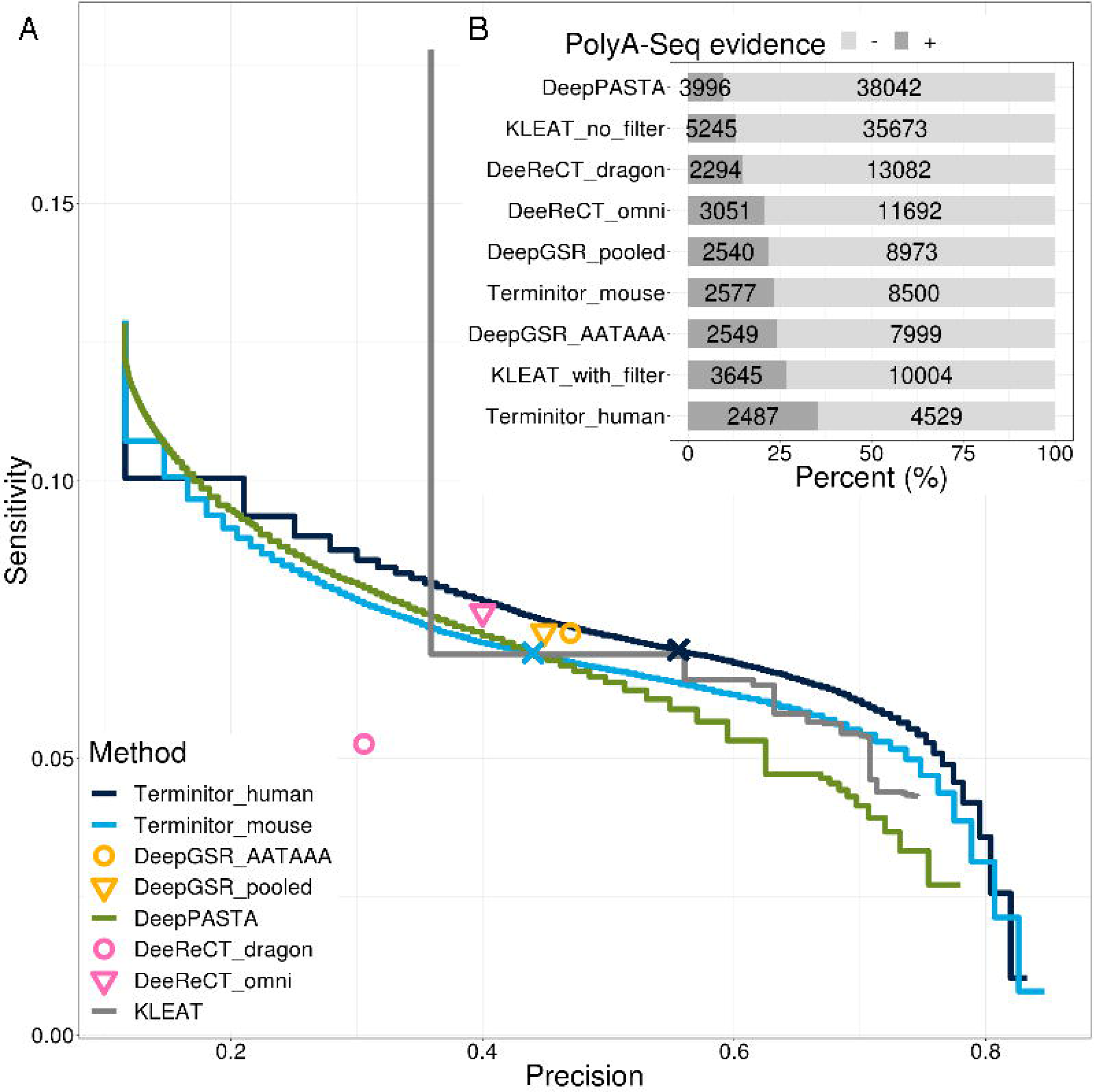
Performance comparison on the UHR sample. **A.** Sensitivity and specificity of poly(A) CS predictions from Terminitor, KLEAT, DeepPASTA, DeeReCT-PolyA and DeepGSR on Ensembl annotated poly(A) CS. The two pre-trained models of DeepGSR are the one with sequences containing only the strongest PAS hexamer AATAAA, and the one with sequences containing 16 hexamer motifs pooled together. The two pre-trained models of DeeReCT-PolyA are derived from the dragon dataset and omni dataset as described (Xia et al., 2019). Two pre-trained Terminitor models are derived from the human and mouse datasets. The navy/blue crosses on Terminitor human/mouse model represent probability = 0.5 cut-off, respectively. **B.** Percentage of poly(A) CS that are missing from Ensembl annotation are supported/not supported by PolyA-Seq. We used probability = 0.5 cut-off for Terminitor and DeepPASTA predictions.

We also note that DeeReCT-PolyA and DeepGSR are binary classifiers, while Terminitor outputs a probability value that can be tuned in favour of either sensitivity or precision. Nevertheless, Terminitor trained on human sequences consistently outperforms all competing tools in both metrics, except for one data point, which corresponds to the raw, unfiltered KLEAT predictions. However, the precision of raw KLEAT predictions is arguably too low (35.90% and 25.52% for UHR and HBR samples, respectively) to be used for high-confidence profiling. Although KLEAT performs comparably to Terminitor on UHR sample, its performance is surprisingly low on HBR sample regardless of what read evidence is used to filter its raw predictions. It is also worth mentioning that the four models built by DeeReCT-PolyA and DeepGSR all have a precision lower than 50%, while precision of the Terminitor human model (at probability = 0.5 cut-off) is 55.59% and 57.72% for UHR and HBR samples, respectively.

Since a given sample may express novel transcripts or transcripts with novel poly(A) CS, we compared the falsely predicted poly(A) CS determined by Ensembl annotation with the corresponding PolyA-seq predictions to identify putative novel events with PolyA-seq evidence (Fig 2B, S5B Fig). Although Terminitor did not predict the highest number of high confidence poly(A) CS overall, it is always the pipeline having the highest percentage (35.45% and 36.95% for UHR and HBR sample, respectively) of predictions that have PolyA-seq evidence support. This suggests its high precision in detecting novel poly(A) CS when compared to the other methods. Although the raw KLEAT predictions identified a large number of poly(A) CS, only a small proportion can be verified by PolyA-seq. We observed that to maintain precision, certain filters have to be applied to make use of KLEAT predictions, while the choice of filter may be highly library-dependent.

We compared the runtime and peak memory usage of all competing tools under the same computing environment (Table 1). All five pipelines used RNA-Bloom [34] as the *de novo* RNA-seq assembler to start with, and the peak memory usage of all pipelines is the same (37.87 GB), contributed by RNA-Bloom. Terminitor, DeepPASTA, DeepGSR, and DeeReCT-PolyA shared the same pipeline to identify candidate sequences, and the only difference is the NN used for sequence classification, which resulted in runtime differences. With a runtime of 7:06:52 hours, DeeReCT-PolyA is the fastest tool, followed by Terminitor with less than 5-minute (1.17%) run time difference. DeepPASTA took two hours more than the first two competitors because of the RNA structure prediction step, although this extra step did not help improve its prediction performance. The long runtime of DeepGSR is probably due to its implementation, which was only compatible with Python 2.7 and older versions of Keras and Theano. KLEAT is more than three times slower compared to all other tools because it involves computationally expensive read-to-contig and contig-to-genome alignments.

**Table 1.** Runtime comparison. The runtime for all four pipelines includes *de novo* assembly of UHR sample (two replicates), alignments, candidate poly(A) CS identification and classification.

| Pipeline | Wallclock Runtime (hh:mm:ss) | Wallclock Runtime without assembly |
| --- | --- | --- |
| DeeReCT-PolyA | 7:06:52 | 17:29 |
| Terminitor | 7:11:23 | 22:00 |
| DeepPASTA | 9:17:49 | 2:28:26 |
| DeepGSR | 11:10:22 | 4:20:59 |
| KLEAT | 25:17:00 | 18:27:37 |

### Impact of expression level on prediction accuracy

The Ensembl annotation is a repertoire of transcript annotations from multiple tissues and cell lines in a range of biological states, so not all of the annotated poly(A) CS are expressed in the HBR and UHR samples, as noted above. We quantified the expression levels of all polyadenylated reference transcripts in Ensembl, and then examined how the four tools perform with respect to poly(A) CS at different expression levels (Fig 3, S6 Fig). Aside from the spikes corresponding to KLEAT raw predictions, Terminitor consistently performs better than all competitors in four tiers of expression thresholds. As expected, the sensitivity of all tools increases as the expression level threshold increases, because unexpressed transcripts are removed from the denominator. The performance difference between Terminitor and other tools increases as well when the expression cut-off increases. In fact, the performance metrics of Terminitor mouse model on poly(A) CS expressed at > 20 TPM are similar to, or better than, DeepPASTA, DeepGSR and DeeReCT-PolyA models for both UHR and HBR samples, showing the promising application of Terminitor on species without a good reference transcriptome annotation.

**Fig 3.**
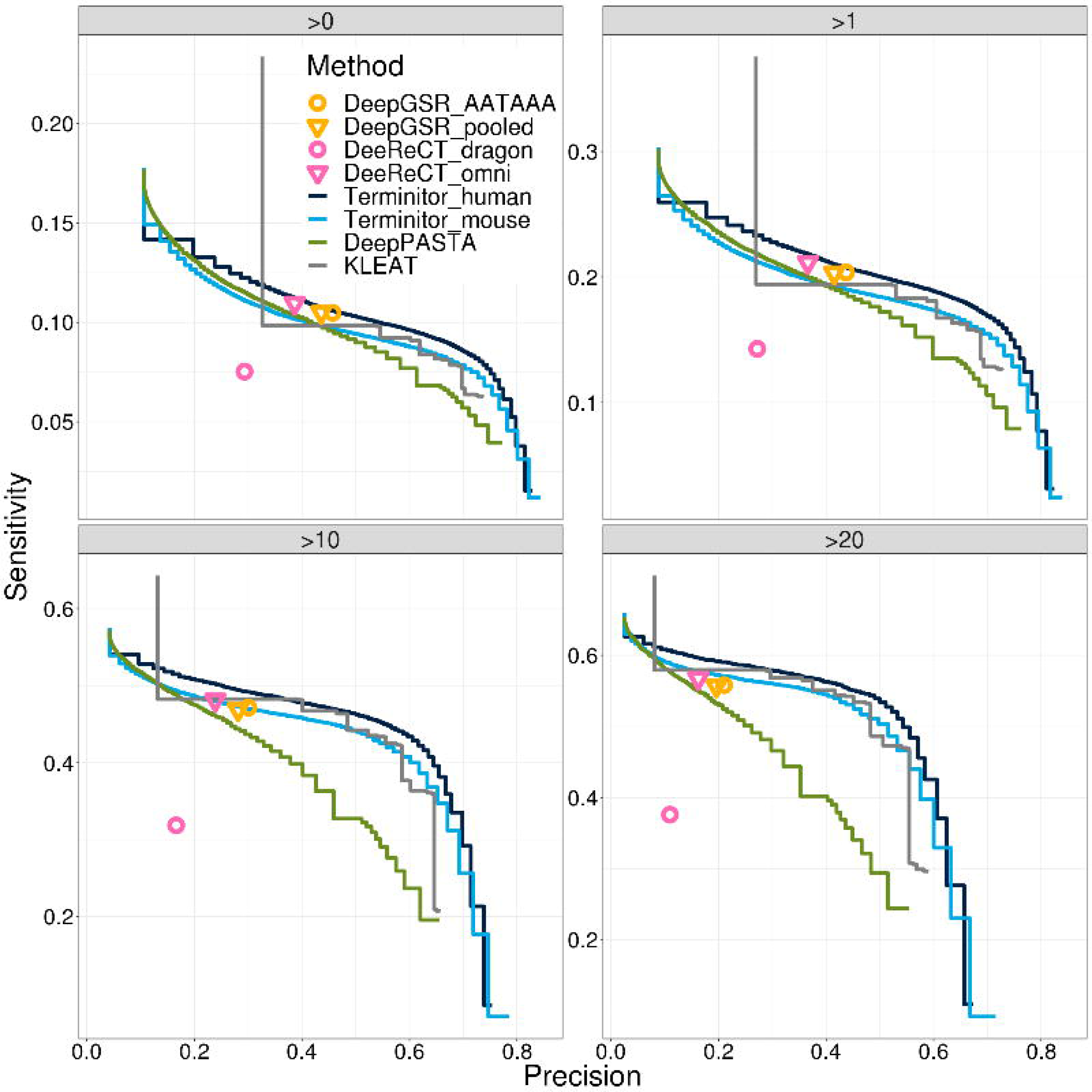
Performance comparison on the UHR sample with different expression level cut-offs. Sensitivity and precision of poly(A) CS predictions of Terminitor, KLEAT, DeepPASTA, DeeReCT-PolyA and DeepGSR on expressed Ensembl transcripts. The four facet plots represent the comparison between all expressed transcripts, expressed transcript with TPM > 1, > 10, and > 20, as indicated.

### Single nucleotide variation in PAS affects APA choices

One of the advantages of Terminitor is that it is capable of predicting poly(A) CS solely based on sequence context. Consequently, poly(A) CS created or destroyed by base changes can also be identified. We carefully examined all candidate sequences belonging to the UHR sample, and identified sequences containing different PAS hexamer motifs from the reference human genome assembly. Through comparisons between the predicted probabilities and the base variation, we discovered two cases of variation-associated polyadenylation.

The first variant base is recorded in dbSNP as rs6484833, a C-to-T change at chr11:36273982 (GRCh38.p12) on the reverse strand as seen in Integrative Genomics Viewer (S7A Fig) [35,36]. This polymorphism creates the strongest hexamer AATAAA on gene *COMMD9*, and it is associated with an unannotated poly(A) CS at chr11:36273960 on the terminus exon. We speculate that this association may be causal, the former causing the latter. In our pipeline, RNA-Bloom successfully reconstructed this novel transcript to full length with a poly(A) tail, and Terminitor predicted it as a poly(A) CS with 99% probability. This poly(A) CS is also detected in PolyA-Seq of the same sample. Based on Ensembl annotation, *COMMD9* has three poly(A) CS associated with the same terminus exon. This site results in the second-longest 3’ UTR, which harbours three miRNA binding sites and extensive RNA binding protein (RBP) sites for 19 RBPs (S8 Fig).

The second variant base is rs15342, a T-to-C change at chr15: 101070089 on gene *LRRK1* (S7B Fig). This polymorphism destroys the strongest hexamer AATAAA, which is associated with an annotated poly(A) CS 22 nt downstream. There are 21 sequencing reads containing this polymorphism extend beyond this poly(A) CS, while reads carrying the wildtype are all polyadenylated. RNA-Bloom successfully assembled both the wildtype transcript with a poly(A) tail and the variant transcript without poly(A) tail. Terminitor classifed the wildtype and variant transcripts to contain poly(A) CS with a probability of 98% and 1%, respectively. This polymorphism may prevent cleavage at the poly(A) CS immediately downstream, leading to the alternative usage of the distal poly(A) CS, and producing a transcript with longer 3’ UTR. Similarly, extensive (16) miRNA binding sites and RBP sites for 77 RPBs are also predicted to exist on this extended 3’ UTR (S9 Fig).

### Terminitor performance on long reads

Although the current short read sequencing followed by *de novo* transcriptome assembly is part of Terminitor’s mainstream RNA-seq analysis pipeline, the advent of long read sequencing technologies provides unprecedented opportunity to study APA at the transcript isoform level. Here we applied the Terminitor pipeline on two sequence libraries from the human sample NA12878, namely Illumina short reads and PacBio circular consensus sequence (CCS) reads (see Methods for details). Terminitor consistently profiles three times or more poly(A) CS in the PacBio library as in the Illumina library, even under different expression level cut-offs (Fig 4). However, with the Illumina library, Terminitor is able to detect more poly(A) CS at a higher precision (> 70%). As the probability cut-off becomes more stringent, less positive poly(A) CS are predicted by both libraries, but the overlapping percentage with respect to the Illumina library increases (S10 Fig). For predicted poly(A) CS with a probability > 90%, 59.27% of the CSs found in the Illumina library are also found in the PacBio library.

**Fig 4.**
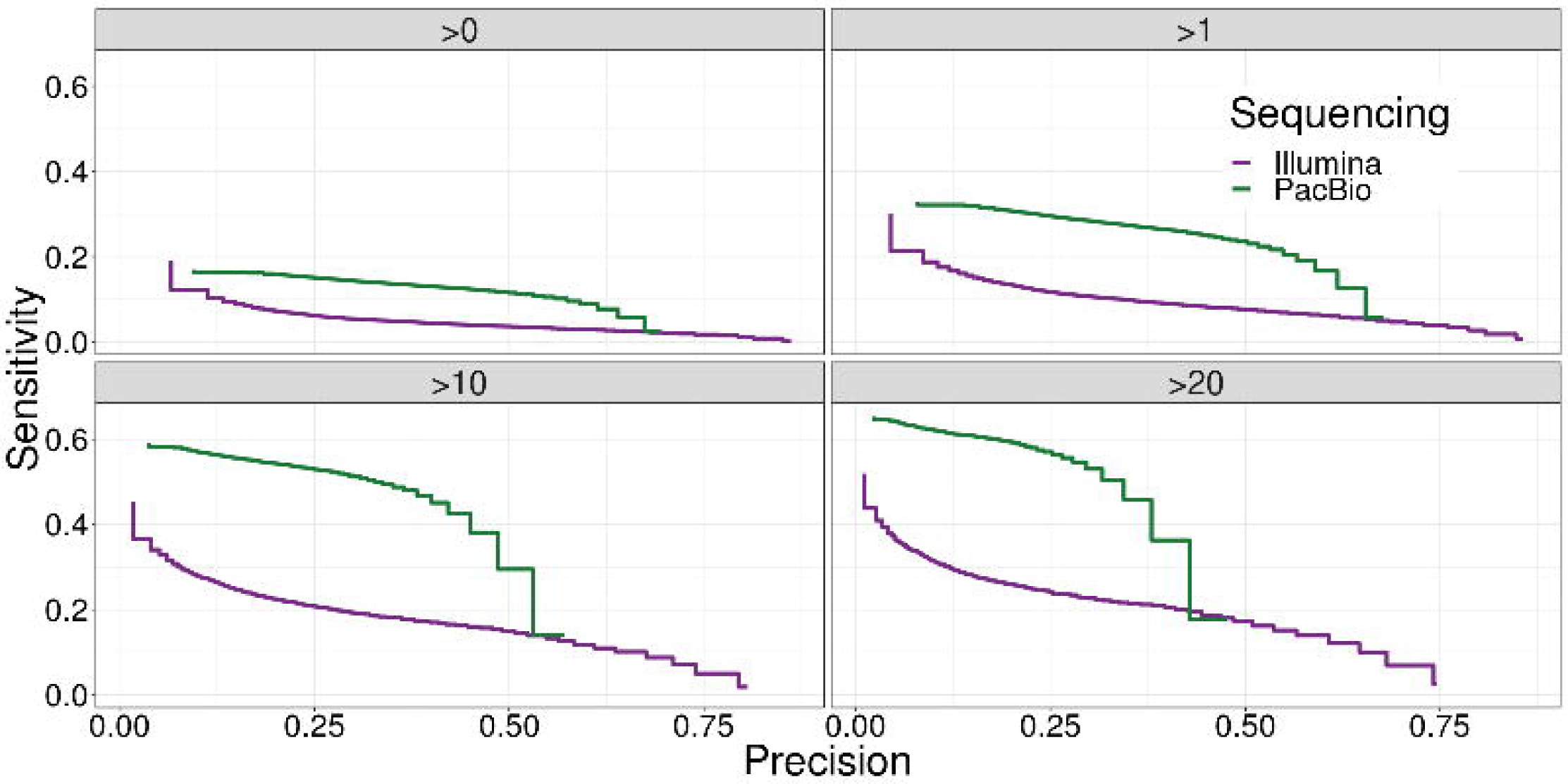
Performance comparison between PacBio library and Illumina library on a *H. sapiens* NA12878 sample. Sensitivity and precision of poly(A) CS predicted by Terminitor on PacBio and Illumina libraries at different expression level cutoffs. The four facet plots represent the comparison between all expressed transcripts, expressed transcript TPM > 1, > 10, and > 20, as indicated.

## Discussion and Conclusions

Here we introduce a novel deep learning model named Terminitor to detect and classify 3’ UTR cleavage sites with or without poly(A) tails, and a profiling pipeline that starts from raw RNA-seq reads to characterize expressed poly(A) CS. The novelties of our model and contributions in this work are as follows:

1. The use of concatenated *k*-mer representations instead of raw sequence as model input. Previous applications of deep learning in omics research have shown that designing a proper representation of the raw data is critical to facilitate classification performance. Inputting additional information, namely well-studied sequence features, can also alleviate the burden of learning complex structures. In the case of polyadenylation, several *cis*-acting elements have been revealed, such as the GU-rich sequences in the downstream, UGUA elements in the upstream, and the conserved hexamer motif. Inspired by the usage of *k*-mer in the genome/transcriptome assembly tasks, Terminitor uses *k*-mer representations of raw sequences to mimic functional motifs. Since the auxiliary elements may have different lengths, multiple *k*-mer representations are jointly used to, and are observed to push the accuracy higher.
2. The use of an embedding layer to learn the similarities between *k*-mer pairs. One-hot encoded *k*-mer representation of raw sequence is high dimensional and sparse. In our case, the input dimension is exponential to *k*, and most positions in the input vector are zeros. This is computationally inefficient, and would not capture similarities between *k*-mers. To address this, we borrowed the concept of word embedding from natural language processing and used it as an interface between the input layer and the subsequent hidden layers. Essentially, in the embedding layer, one-hot encoded *k*-mers are converted into dense vectors, representing the projection of *k*-mers in a continuous vector space. Consequently, motifs with similar functions are likely to have similar weights, and treated similarly in the subsequent layers.
3. Separation of non-polyadenylated CSs from polyadenylated CSs. To our knowledge, Terminitor is the first poly(A) CS classifier that includes non-polyadenylated CS label. Although most protein-coding and long-noncoding RNAs contain poly(A) tails at their 3’ ends, a significant number of functional transcripts are not polyadenylated [37]. Since the 3’ end processing mechanism of this group is distinct from polyadenylated CS group, separating it out as a new class improves the performance of the model. Sequence clustering based on Terminitor’s last hidden layer also groups sequences containing non-polyadenylated CS and non-CS together as a big cluster (S3 Fig).

Compared to targeted 3’ end sequencing technologies, RNA-seq data, whether generated using short or long read techniques, are more accessible and widely used in transcriptome studies. In the reported work, we demonstrate that the Terminitor pipeline can identify poly(A) CS on expressed transcripts with high precision in comparison to the state-of-the-art methods for RNA-seq data. Failure in detecting a real and expressed poly(A) CS is mostly due to the failure of reconstructing transcripts to their 3’ ends. For instance, in the UHR sample, among the 58,925 expressed poly(A) CS that were not detected, more than half (30,181) have a TPM less than one, and the majority of them have very few or no reads supporting the poly(A) tail because of sequencing bias.

We observed that false positive predictions by Terminitor generally occur for three reasons. The first and most common scenario is mis-assemblies due to the low sequence complexity of the 3’ UTR. Second, these poly(A) CS are in fact real novel events, but unannotated; around 30% of the Terminitor false positives (using the Ensembl annotation) are corroborated by PolyA-Seq evidence (Fig 2B, S5B Fig). Third, mis-alignment of upstream sequences, meaning the sequence may contain a real poly(A) CS, but the genomic location is incorrect. In the Terminitor pipeline, when an upstream sequence is multi-mapped, only the primary alignment is chosen to help reduce the false positive rate. However, when the sequence comes from gene paralogs, the genomic location inferred from the chosen alignment does not necessarily correspond to that of the expressed paralog.

In principle, the Terminitor pipeline can be applied on any reconstructed transcripts to determine if its 3’-end contains a poly(A) CS, or non-polyadenylated CS, or no site (3’ incomplete). Aside from benchmarking its performance on traditional Illumina short read assemblies, we also explored its application on PacBio long read technology. Terminitor demonstrated more than twice the CS prediction sensitivity when using the PacBio vs. Illumina library, probably because more sequences containing real poly(A) CS are fed into the NN model. PacBio CCS reads are self-corrected through consensus calling step, resulting in near- or full-length transcripts with poly(A) tails. In contrast, transcripts reconstructed from an Illumina library may suffer from mis-assemblies, and lowly expressed poly(A) CS may be obscured by longer transcripts. In our analysis, among the 548,448 candidate sequences extracted from the PacBio library, 29.71% of them contain untemplated poly(A)s, while the ratio is much lower (7.22%) in 381,934 assembled sequences using the Illumina library.

In summary, Terminitor outperforms all state-of-the-art methods in the datasets we benchmarked them on, and executes within a reasonable time, making it an attractive tool to complement other RNA-seq analysis pipelines. The performance of Terminitor is robust on both short and long-read sequencing technologies. As long-read RNA-seq finds broader uptake in transcriptome studies, it is poised to shed light on the association between poly(A) CS and transcript isoforms, fueling a new era of APA research. Terminitor is a candidate enabling technology for such studies. Further, we have observed how Terminitor can identify novel poly(A) CS caused by base variations. As has been elucidated in previous studies, novel poly(A) CS are expressed in cancer tissues, and with the help of Terminitor, mechanisms and impacts of APA can be better understood. Even for species without an established poly(A) CS database, their expressed poly(A) CS can be identified with Terminitor using transfer learning, where the model is trained on a different but related species. The usage of Terminitor, as a sequence classifier, is not limited to poly(A) CS profiling, but can also be extended to facilitate gene annotation and testing the completeness of assembled transcripts. We expect Terminitor to have broad applications in genome/transcriptome annotation, novel isoform identification, APA analysis and gene regulation studies.

## Methods

### Datasets

To train and validate the NN model, and to demonstrate transfer learning, we collated two reference datasets describing human and mouse transcriptomes, based on poly(A) CS databases and Ensembl annotations. Each dataset is composed of sequences with equal lengths coming from three classes: poly(A) CS, non-poly(A) CS, and non-CS (Table 2). PolyA_DB version3.1 was used as our primary poly(A) CS library [13]. We first mapped poly(A) CS recorded in PolyA_DB3 to Ensembl annotation release 94 using `bedtools closest` (v2.27.1) and selected the most compatible transcript for each site. Poly(A) CS supported by less than 1 Reads Per Million (RPM) or located more than 1000 nt downstream of any annotated transcript were considered unreliable and were discarded.

**Table 2.** Size and origin of datasets for model training, testing, and validation.

| Species | poly(A) CS | non-poly(A) CS | non-CS |
| --- | --- | --- | --- |
| *H. sapiens* | 52,457 | 27,595 | 52,460 |
| *M. musculus* | 75,661 | 37,028 | 75,660 |

To avoid potential poly(A) CS contaminating the other two classes, we expanded our poly(A) CS library by incorporating three additional poly(A) databases, including APADB v2, APASdb, and PolyASite. Again, we used `bedtools closest` to assign poly(A) CS in the compiled poly(A) library to Ensembl transcripts. Transcripts with no poly(A) CS from the above-mentioned databases assigned were considered as cleaved but not polyadenylated, and their 3’ ends constituted the non-polyadenylated CS set.

Non-CS comprises five types of genomic locations: exonic, intronic, intergenic, 5’ and 3’ untranslated regions, and are randomly selected at least 100 nt away from any annotated 3’ ends, including Ensembl annotation and compiled poly(A) library.

Our datasets are imbalanced in their label abundances due to the limited number of non-polyadenylated CS. However, to fully exploit the potential of deep NNs, we used all non-redundant poly(A) CS and CS generated as above descriptions, and deliberately matched the number of pseudo sites to the number of poly(A) CS. Because the label abundances are imbalanced, we used both accuracy and the F-measure, the harmonic mean of precision and recall, as assessment metrics during model training and benchmarking.

### Model training

The NN model is implemented in Python 3.6.6, using Keras library 2.2.4 [38] with TensorFlow 1.11.0 [39] as the backend. To evaluate the performance of the model with different hyperparameter combinations and input sequence lengths, we used stratified 5-fold cross validation on the whole dataset to monitor the overall accuracy of three labels, and accuracy, sensitivity, and specificity on poly(A) label. All performance metrics on the training sets reported in this study are the mean and standard deviation of five validation runs. In each run, 80% of data is used for training and 20% is used for validation. Next, we trained the model using the entire training set and used it for benchmarking the performance on experimental data.

### Model architecture and hyperparameter tuning

Searching for the optimal hyperparameters for a deep NN has always been a tedious task, as many parameters are dependent on each other in the way they influence performance. We first built the basic frame with one input layer, one embedding layer, and one output classification layer, and then fine-tune hyperparameters related to architecture and training using grid search, as listed in Table 3. For all models, initial model weights were randomly drawn from Glorot uniform initializer [40]. The weights of the three neurons in the output layer are computed via softmax activation function, representing the probability distribution of the three labels, and they always add up to one. To avoid over-fitting, we applied early stopping in training each model. As an iterative method, the validation loss is monitored at each training epoch, and when it stays the same or increases for the next ten epochs, the training process is stopped. Without early stopping, too many training epochs will lead to nearly 100% accuracy on training set, yet yield very low accuracy on validation and test sets.

**Table 3.** Model hyperparameter tuning. Bolded parameters are the selected optimal values

| Parameter type | Parameter | Search space |
| --- | --- | --- |
| Training | Optimizer | SGD, **Adam**, Adagrad, RMSprop |
|  | Learning rate | 0.1, 0.01, **0.001**, 0.0001 |
|  | Loss function | **Categorical_cross_entropy**, mean_squared_error |
|  | *K*-mer representation | {1}, {2}, {3}, {4}, {5}, {6}, {7}, {8}, {9}, {10}, {11}, {12}, {4, 5}, {4, 6}, {4, 8}, {4, 10}, {4, 11}, {4, 5, 6}, {4, 6, 8}, {4, 5, 6, 10}, **{4, 6, 8, 10}**, {4, 5, 6, 8, 10} |
|  | Batch size | 32, **64**, 128, 256 |
| Architecture | Number of layers | 1, 2, 3, **4**, 5 |
|  | Number of nodes for layers | 4, 16, **32**, **64**, **128**, 256, **512**, 1024<br>(Refer to Fig 6 for details) |
|  | Regularization | Dropout: **0**, 0.1, 0.2, 0.5 |
|  | Activation function for hidden layers | tanh, **ReLU** |
|  | Activation function for output layer | **Softmax**, Sigmoid |

The final architecture of Terminitor is depicted in Fig 5 and the selected optimal hyperparameters are in bold font in Table 3. The first step of Terminitor is to concatenate all *k*-mer representations of the raw DNA sequence. Each *k*-mer in the one-hot encoded raw sequence is represented as a vector of length *l*, where

**Fig 5.**
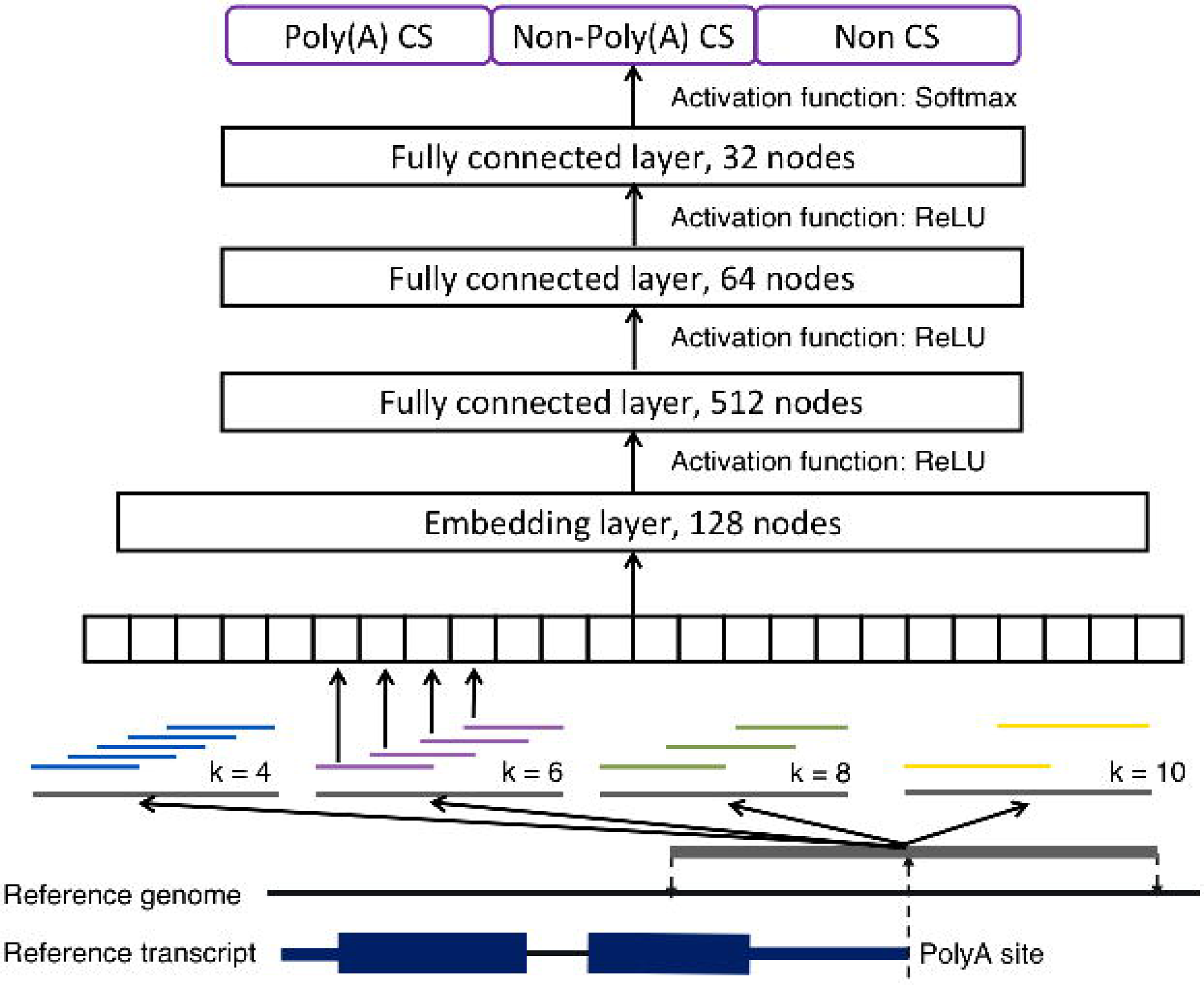
Architecture of Terminitor. The information flows from bottom up. During training stage, reference transcripts are aligned back to the reference genome, and flanking sequence on both side of the poly(A) CS is extracted (grey thick sequence). The sequence is *k*-merized and four *k* values are used. Then all *k*-mer representations are concatenated and each *k*-mer is one-hot encoded (each square represents one *k*-mer). The input sequences are fed into an embedding layer, followed by three fully connected layers, and then the probability of belonging to one of the three classes is calculated through a Softmax function.

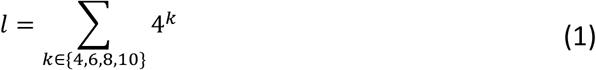

Then an embedding layer with 128 nodes is used to connect the input layer and hidden layers. The next four layers are all fully connected, and the activation function between each adjacent layer is the rectified linear unit (ReLU) function. Since dropout regularization did not achieve better performance, it was not performed in the final model.

### Experimental data

The raw RNA-seq reads for the two MicroArray Quality Control project reference human samples, HBR and UHR, were downloaded from Illumina BaseSpace, and each sample has two replicates (https://basespace.illumina.com/projects/3777775/samples) [41]. RNA-seq libraries of the four samples were prepared with Illumina TruSeq Stranded kits using poly(A) selection, and sequenced as 75bp paired-end reads. The Illumina and PacBio reads for NA12878 were obtained from an earlier study under accession number SRR1153470 and SRR1163655, respectively [42]. The Illumina reads are from a poly(A) selected library sequenced with 75 bp stranded paired-end reads. The PacBio library was prepared from the same batch of poly(A) selected RNA and sequenced with PacBio CCS method. The filtered PolyA-Seq sites were downloaded from the NCBI Gene Expression Omnibus (GEO) under study GSE30198. We combined sites detected by the two HBR replicates (GSM747473 and GSM GSM747474), and sites detected by the two UHR replicates (GSM747475 and GSM747476) together respectively for benchmarking analysis.

### Prediction pipeline

We also provide a poly(A) CS prediction pipeline based on Terminitor. When using Illumina short reads, the pipeline starts by assembling RNA-seq data using RNA-Bloom [34]. RNA-Boom is run with the option `-stratum 01`, which allows extension of all fragments regardless of its coverage, and the option `--polya`, which prioritizes the assembly of transcripts with poly(A) tails. The false positive rate (FPR) of Bloom filters in the program is set to 0.005. Then, all assembled transcripts are aligned to the reference genome hg38 and compared to Ensembl annotation release 94 to select sequences with untemplated As at their 3’ ends, or sequences that end in the 3’ UTR of annotated transcripts. Similar to the structure of sequences in the training dataset, selected sequences are used to construct test sequences, each composed of three parts: a fixed length (100 nt) of upstream sequence extracted from the assembled transcript, a potential cleavage site, and a fixed length (100 nt) of downstream genomic sequence extracted from the reference. Finally, these test sequences are fed into the trained Terminitor model for classification.

### Evaluation of prediction pipeline

Poly(A) CS are often observed as a group of neighboring sites rather than a precise genomic location [7]. To accommodate, predicted poly(A) CS within 30 nt of each other are clustered, and the one with the highest probability score is chosen. Ensembl annotated poly(A) CS are also clustered in the same way, and the one located in the median position is chosen as the representative. Then, the predicted list is compared with the annotated list to compute hits and misses. In our evaluation system, a true positive poly(A) CS is a predicted one that lies within ± 30 nt of an annotated site; a false positive site is a predicted poly(A) CS that does not lie within ± 30 nt of any annotated ones; false negative means an annotated site does not have any predicted site within ± 30 nt. For this analysis, true negative (no real or predicted poly(A) CS in a certain location) is not informative for experimental data. Using these definitions, we report sensitivity and precision as performance measures for all experimental data tests. Note that in the training data, true negative is informative, and we also report accuracy for our cross validation results.

### Tool comparison

We applied the proposed prediction pipeline on experimental data and compared its performance with the performance of the state-of-the-art methods. All experiments were run on Centos 6.7 system with 12 Intel Xeon E5-2650 CPUs and 84 GB memory.

KLEAT pipeline was chosen as a representative of read-evidence based approaches. The auxiliary files KLEAT uses include: 1) RNA-seq assemblies from RNA-Bloom, 2) read to contig alignment files generated by BWA-MEM 0.7.17-r1188 [43], and 3) contig to genome alignment files generated by GMAP version 2017-01-14 [44]. Since KLEAT only works for the human genome reference hg19, all predicted poly(A) CS were converted into hg38 genomic coordinates by UCSC liftOver [45]. KLEAT outputs all possible poly(A) CS and their supporting evidence, such as the length of poly(A) tail in reconstructed transcripts, the number of reads containing poly(A) tail, and the length of poly(A) tail of reads that can be mapped to a reconstructed transcript. We applied filters to the raw predictions with different combinations of three evidence types and computed the Pareto frontier on the sensitivity-precision plane.

Two recent deep learning models, DeepGSR and DeeReCT-PolyA, were also included in the comparison. Like Terminitor, these two tools are NN models expecting fixed length sequences as input. Starting from all candidate sequences processed from our prediction pipeline, we selected only the ones containing PAS as these tools expects PAS in the input sequences. Then we adjusted the sequence lengths as required by the two models (± 300 nt around the PAS for DeepGSR; ± 100 nt around the PAS for DeeReCT-PolyA), and fed into these two models for classification. For DeepGSR, we used two pre-trained models from its publication, one only on the AATAAA hexamer, and the other on all 16 hexamers [29]. Similarly, the DeeReCT-PolyA package provides two pre-trained models based on two human datasets, dragon and omni, both of which were included in the comparison. Both datasets were generated by previous poly(A) prediction researches [46,47]. Both DeepGSR and DeeReCT-PolyA only output a binary value instead of a confidence score that can be tuned with different cut-offs, so their predictions contributed to four data points in the comparison plot: two models for two tools.

### Data access

Terminitor is implemented using Python 3.6 with Keras package. The software package is freely available under GNU General Public Licence. The source code and manual are available at our GitHub repository https://github.com/bcgsc/Terminitor. Previously published data used in this work are described in the Methods section, and are available from several public repositories. The human and mouse reference datasets colated in this study and trained models are deposited in public ftp site http://www.bcgsc.ca/downloads/supplementary/Terminitor.

## Supporting information

**S1 File. Supplementary material.** This file contains all supplementary figures S1-10 Fig.

