## Supplementary material for "Terminitor: Cleavage Site Prediction Using Deep Learning Models"

### Supplementary Figures

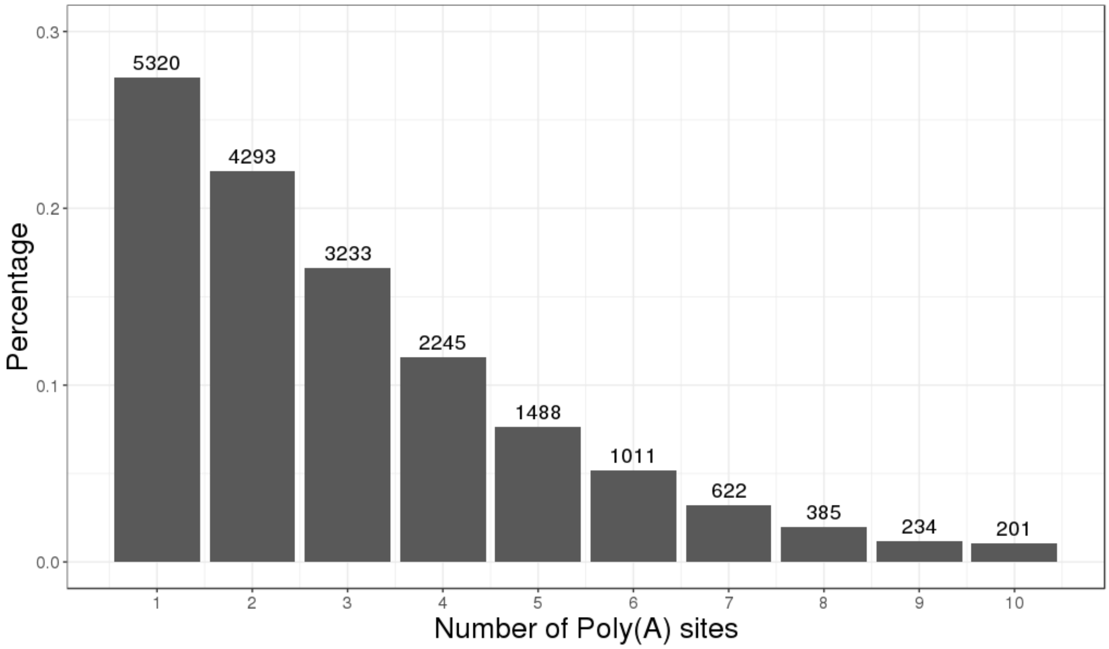


S1 Fig. **Histogram of APA isoforms in Ensembl annotation GRCh38.94.** Poly(A) sites more than 10 are not shown here, and they constitute 2.1% of genes.

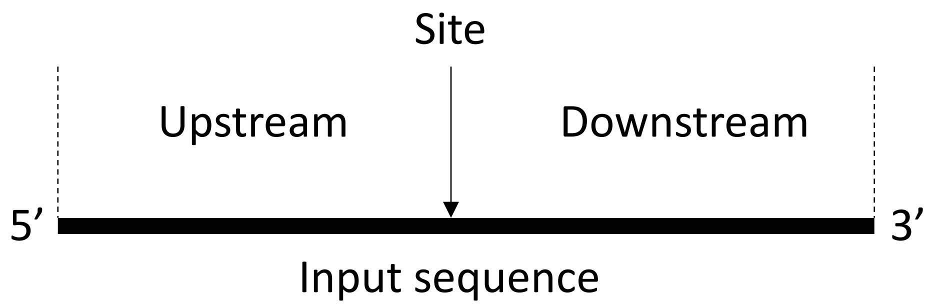


S2 Fig. **Schematic representation of the input sequence.**

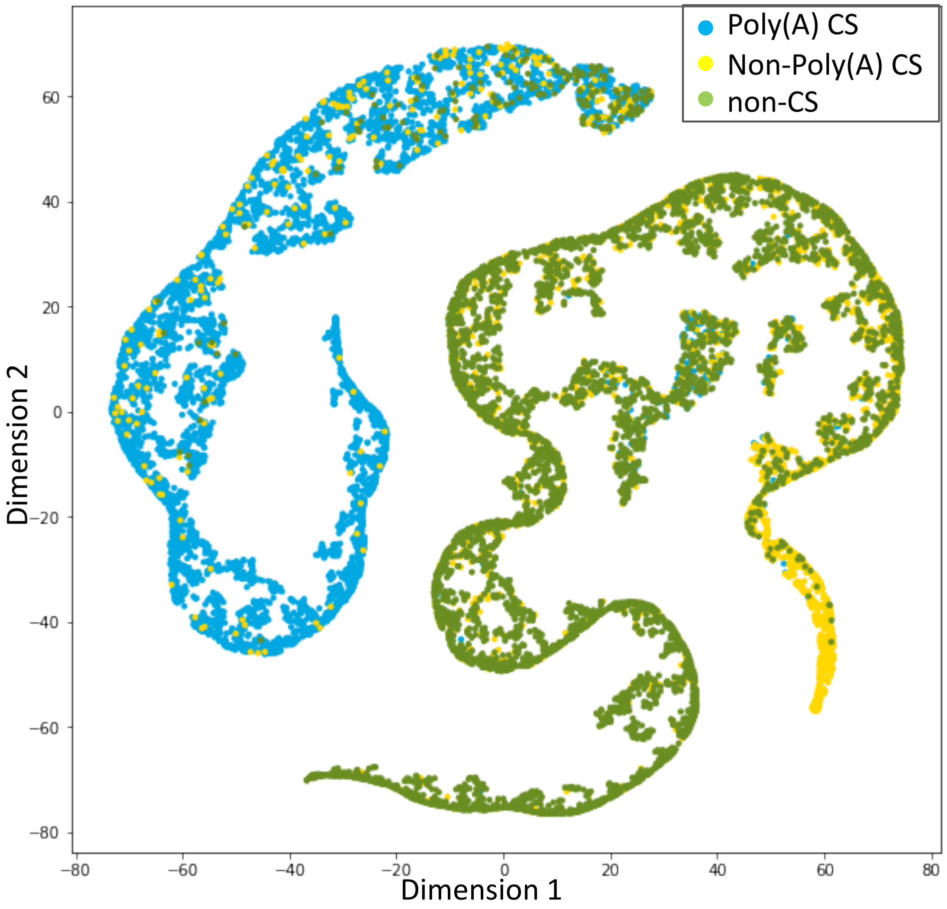


S3 Fig. **T-SNE plot visualizing the separation of three sequence classes.** Each dot represents a test sequence of length 200 nt (100 nt upstream + 100 nt downstream). Sequences are projected into t-SNE space based on the weights of the last hidden layer from Terminitor, with the first two components plotted as the axes of the plot. Cluster assignments of sequences are based on their real class.

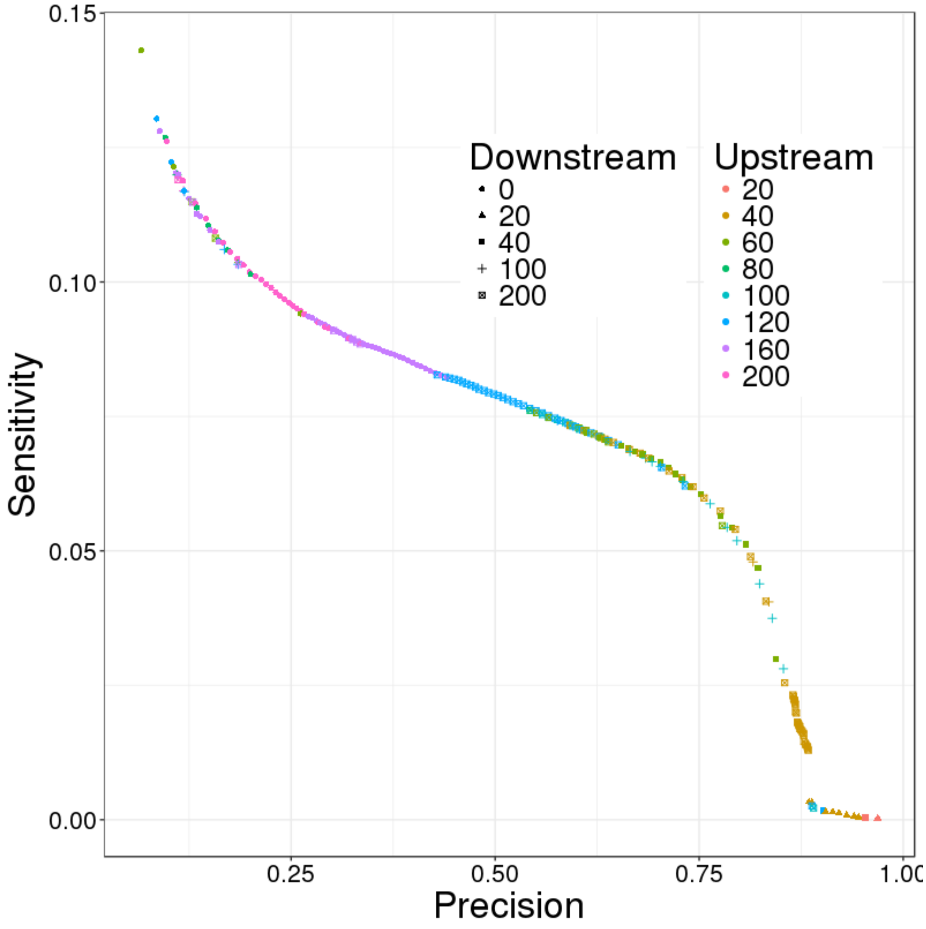


S4 Fig. **Performance of 40 models with different length combinations on the UHR sample.** Only the models whose performance lie on the Pareto Frontier are plotted in the figure. Each model was trained with the same set of sequences but of different up/downstream lengths, and was applied on the same set of candidate sequences extracted from the UHR sample.

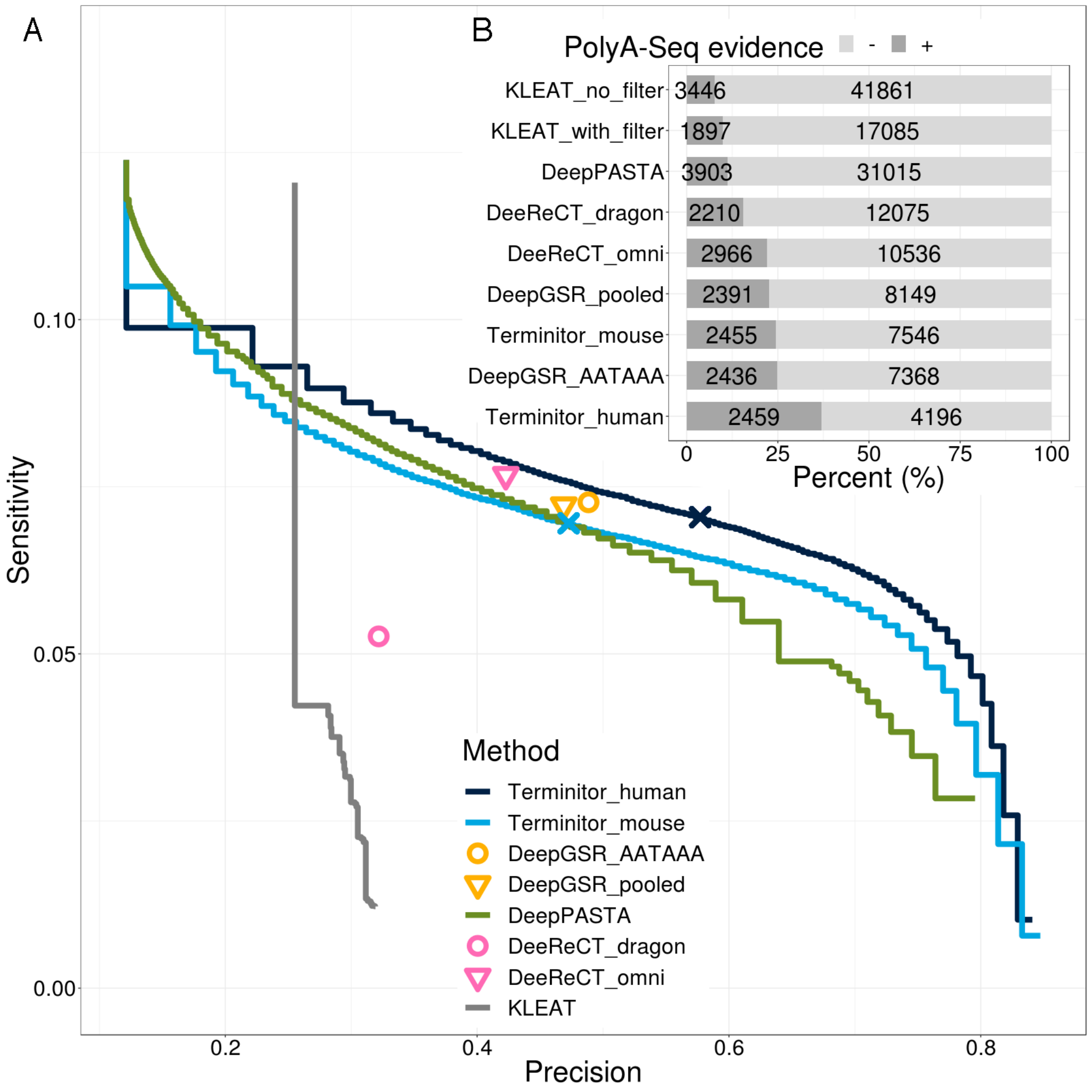


S5 Fig. **Performance comparison on the HBR sample.** A. Sensitivity and specificity of poly(A) CS predictions from Terminitor, KLEAT, DeepPASTA, DeeReCT-PolyA and DeepGSR on Ensembl annotated poly(A) CS. The two pre-trained models of DeepGSR are the one with sequences containing only the strongest PAS hexamer AATAAA, and the one with sequences containing 16 hexamer motifs pooled together. The two pre-trained models of DeeReCT-PolyA are derived from the dragon dataset and omni dataset as described in Xia et al., 2019. Two pre-trained Terminitor models are derived from the human and mouse datasets. The navy/blue crosses on Terminitor human/mouse model represent probability = 0.5 cut-off, respectively. B. Percentage of poly(A) CS that are missing from Ensembl annotation are supported/not supported by PolyA-Seq. We used probability = 0.5 cut-off for Terminitor and DeepPASTA predictions.

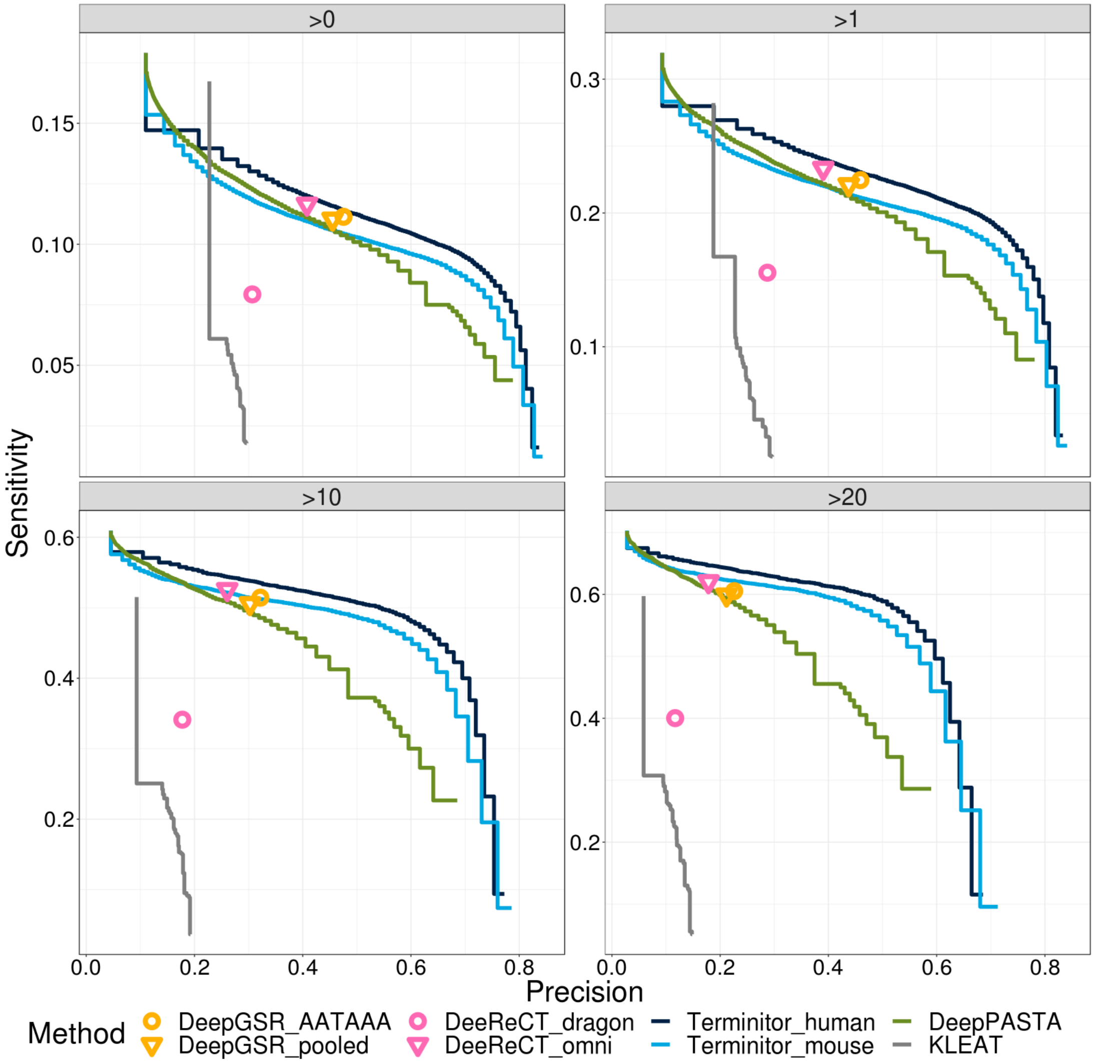


S6 Fig. **Performance comparison on the HBR sample with different expression level cutoffs.** Sensitivity and precision of poly(A) CS predictions of Terminitor, KLEAT, DeepPASTA, DeeReCT-PolyA and DeepGSR on expressed Ensembl transcripts. The four facet plots represent the comparison between all expressed transcripts, expressed transcript with transcript per million (TPM) > 1, > 10, and > 20, as indicated.

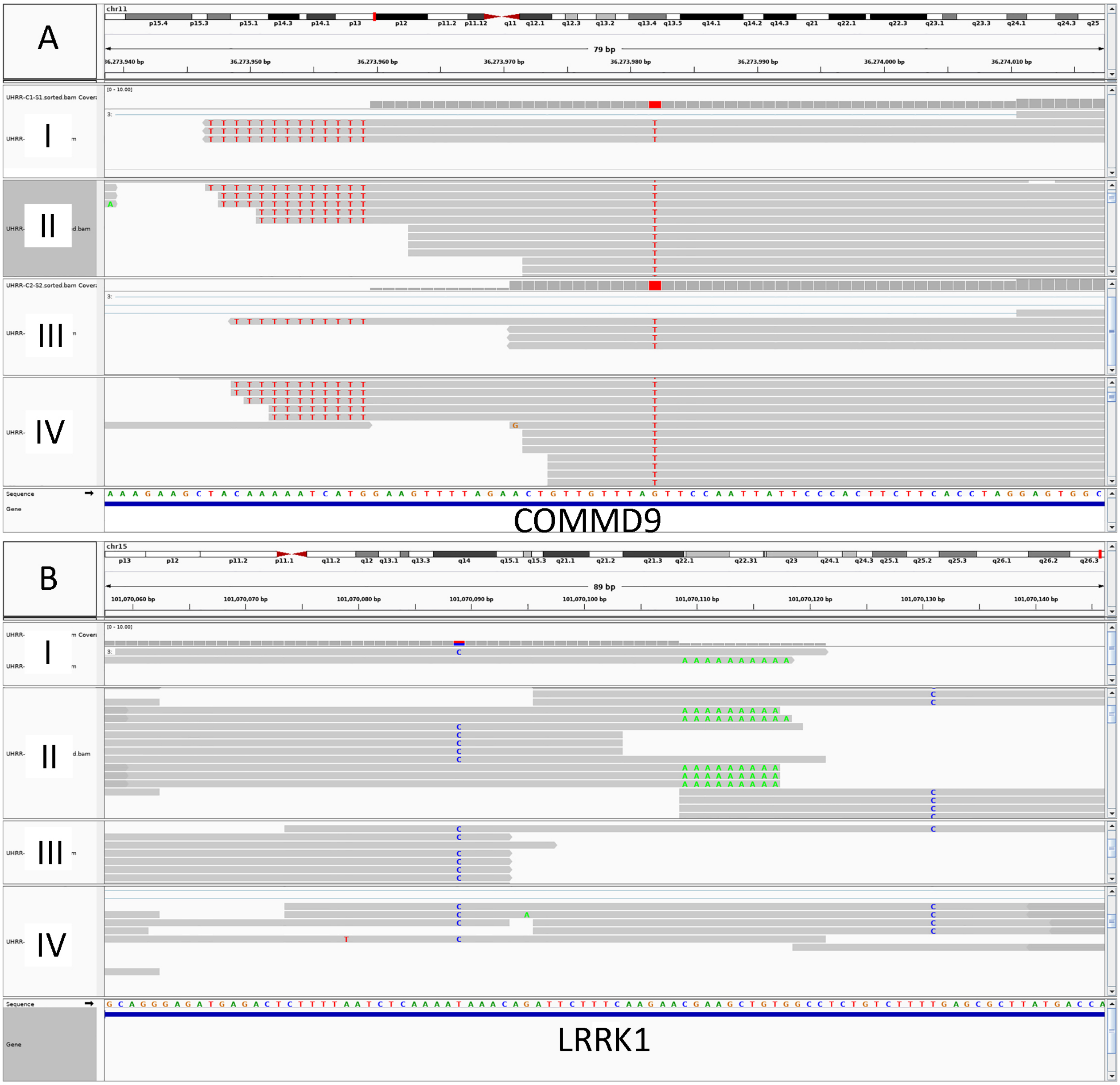


S7 Fig. **Integrative Genomics Viewer screenshot of two SNPs and the corresponding poly(A) CS.** All data shown here are based on the UHR sample. **A.** SNP rs6484833 on gene *COMMD9*, **B.** SNP rs15342 on gene *LRRK1*. For both panels, four tracks from top to bottom are **I**, replicate 1 assembled transcripts; **II**, replicate 1 raw reads; **III**, replicate 2 assembled transcripts; IV, replicate 2 raw reads.

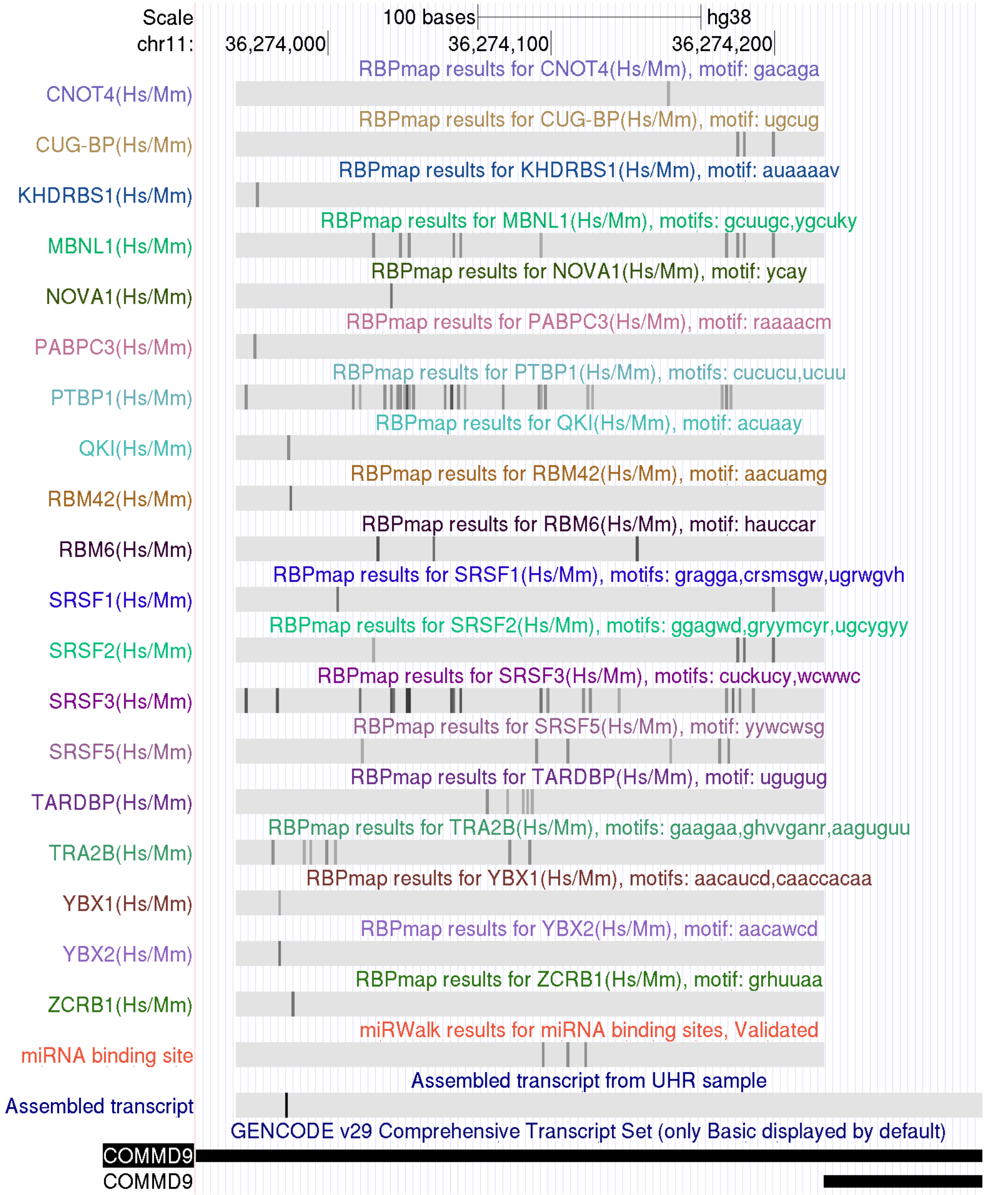


S8 Fig. **MiRNA binding sites and RNA binding protein (RBP) sites on the 3' UTR of COMMD9 with respect to the newly discovered poly(A) CS.** The GENCODE track shows two annotated transcripts with different poly(A) CSs and the assembled transcript track shows the assembled transcript till it's 3' end. The black tick indicates the C-to-A mutation. In the miRNA binding site track, 3 miRNA binding sites validated by MiRTarBase are shown from the end of the shorter poly(A) CS till the end of the newly discovered one. All the rest tracks are RBP binding sites predicted by RBPmap.

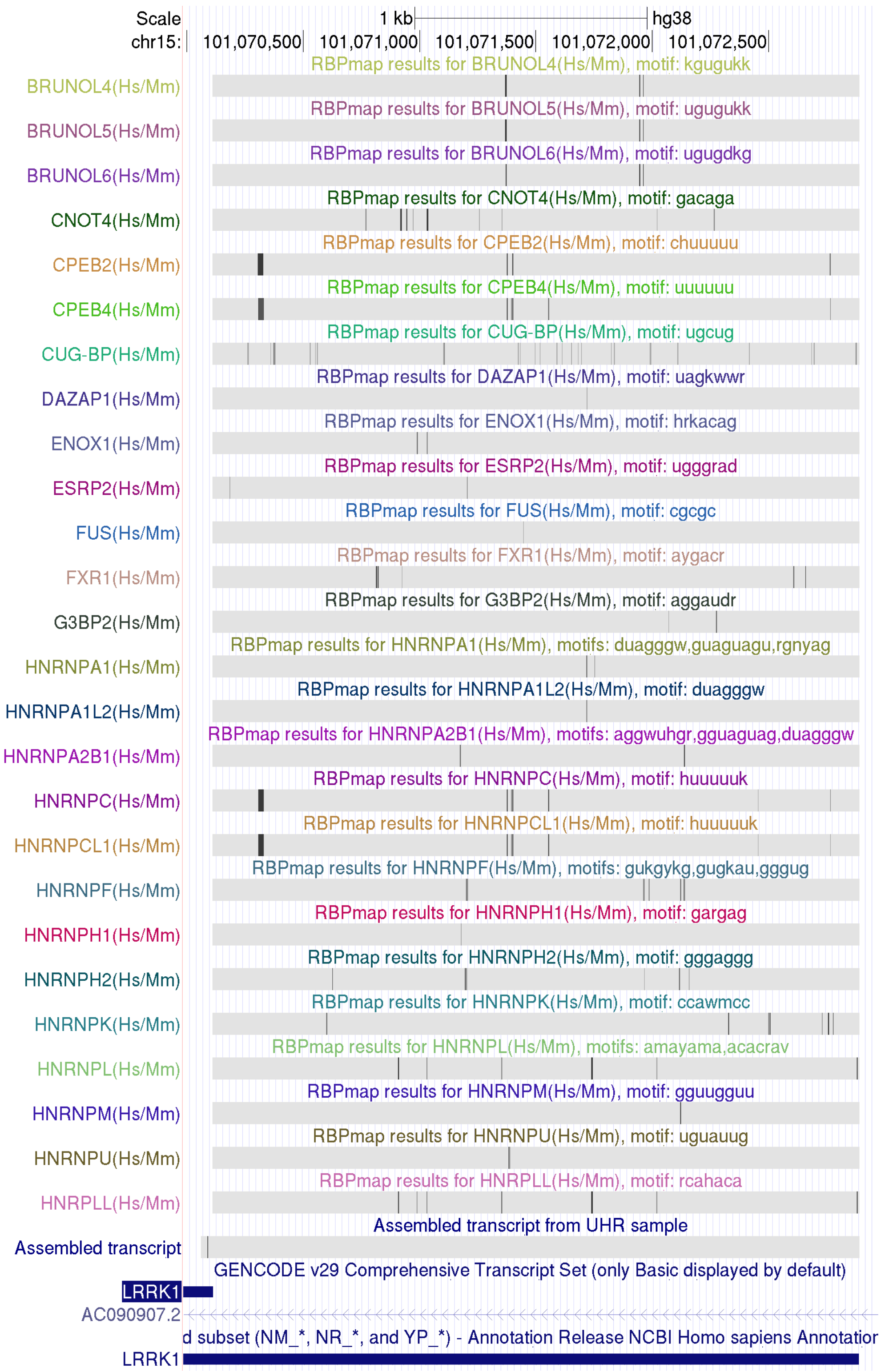


S9A Fig. **RBP sites on the extended 3' UTR of LRRK1 with respect to the destroyed poly(A) CS.** The GENCODE track and RefSeq track show two annotated transcripts with different poly(A) CSs and the assembled transcript track shows the assembled transcript with the C-to-A mutation as the black tick. All the rest tracks are RBP binding sites predicted by RBPmap.

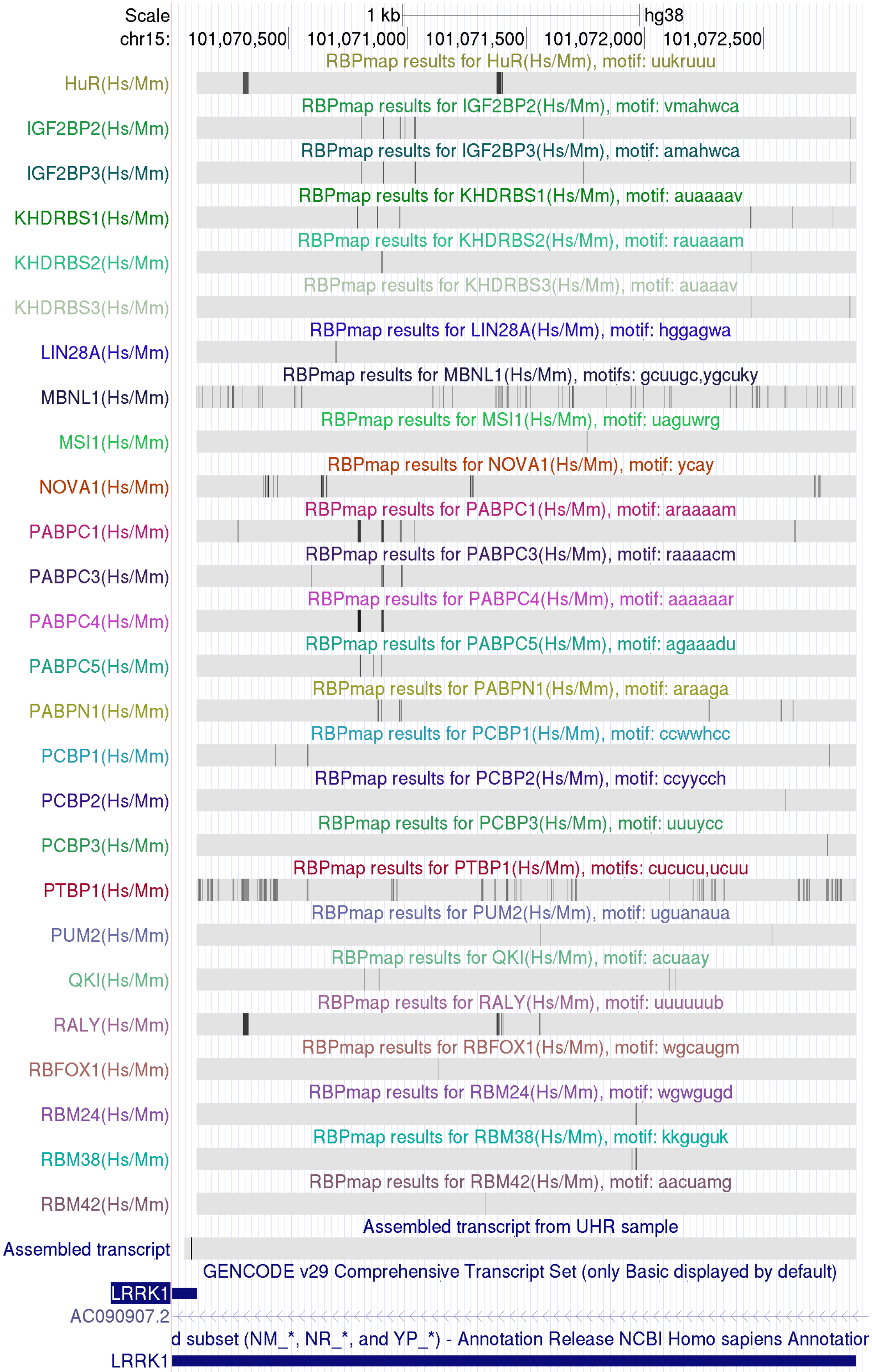


S9B Fig. **RBP sites on the extended 3' UTR of LRRK1 with respect to the destroyed poly(A) CS.** The GENCODE track and RefSeq track show two annotated transcripts with different poly(A) CSs and the assembled transcript track shows the assembled transcript with the C-to-A mutation as the black tick. All the rest tracks are RBP binding sites predicted by RBPmap.

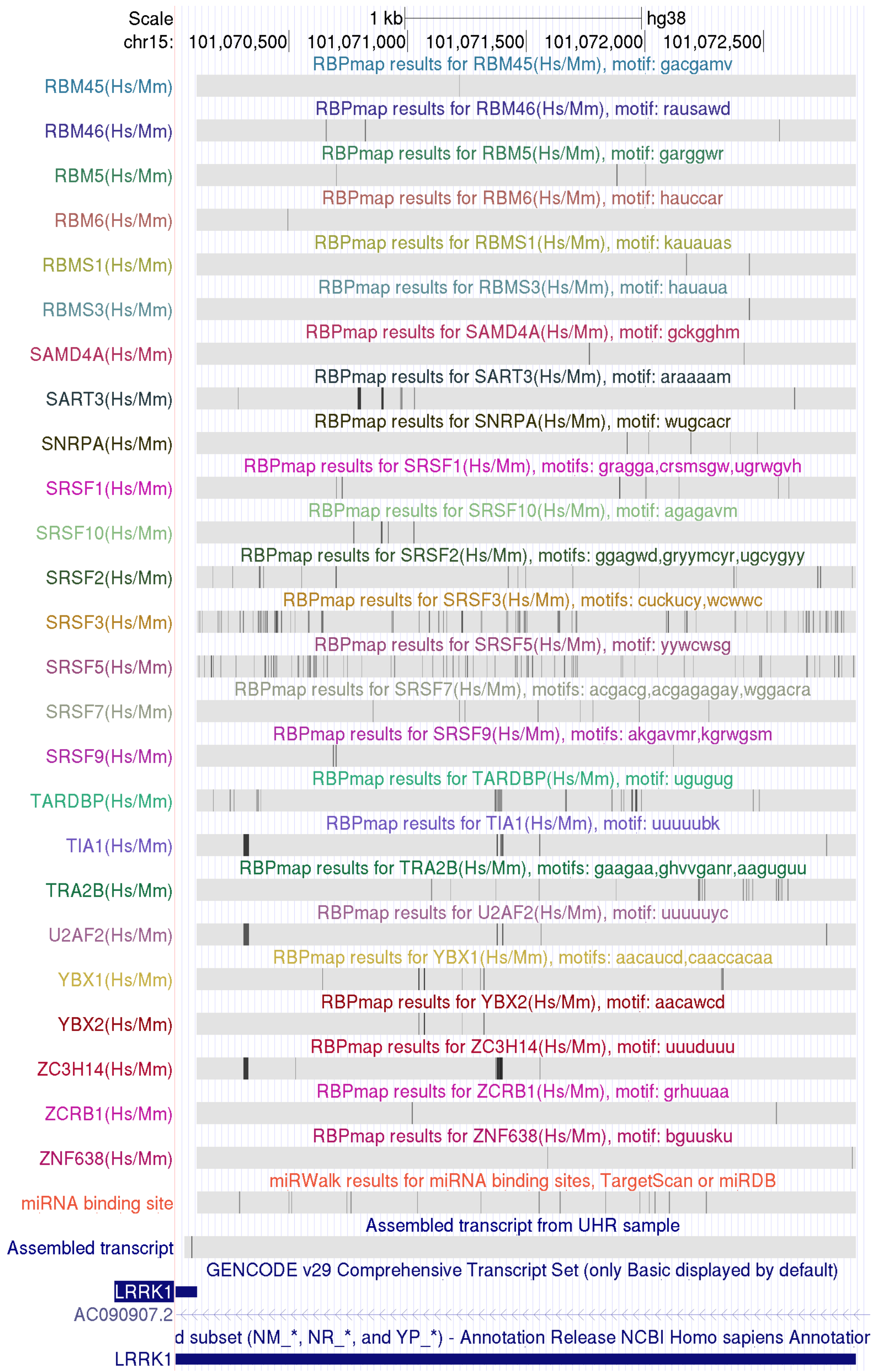


S9C Fig. **MiRNA binding sites and RBP sites on the extended 3' UTR of LRRK1 with respect to the destroyed poly(A) CS.** The GENCODE track and RefSeq track show two annotated transcripts with different poly(A) CSs and the assembled transcript track shows the assembled transcript with the C to A mutation as the black tick. The miRNA binding site track shows all the miRNA binding sites predicted by TargetScan or miRDB. All the rest tracks are RBP binding sites predicted by RBPmap.

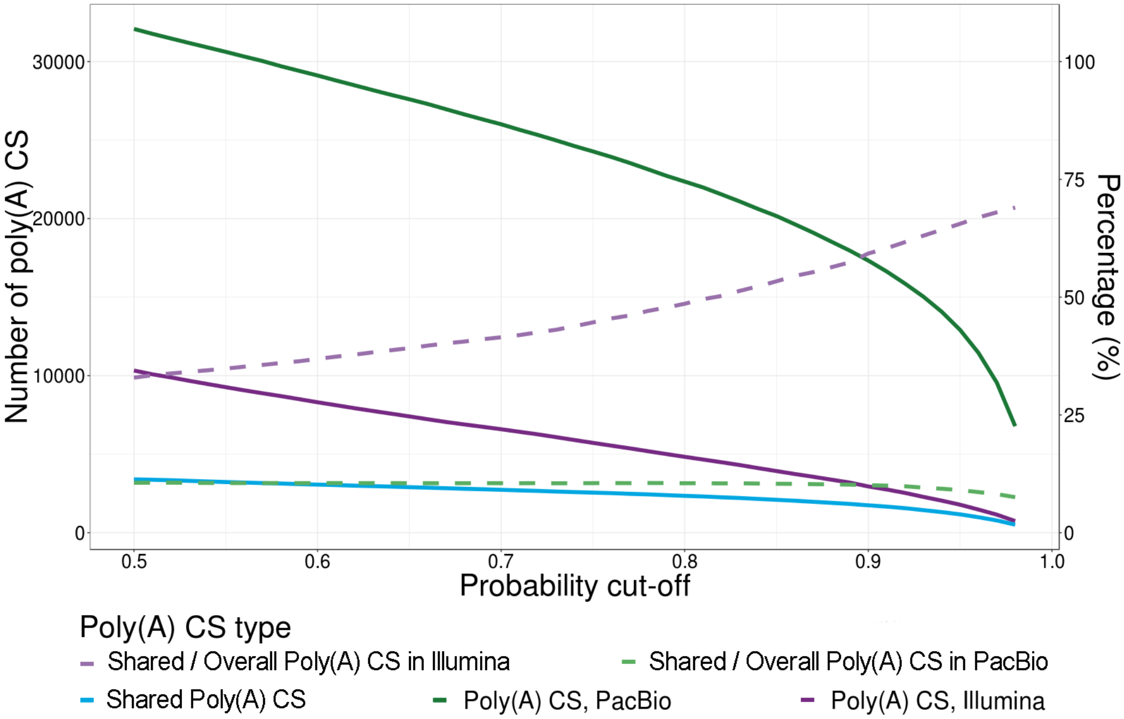


S10 Fig. **Shared poly(A) CSs identified in PacBio long-read and Illumina short-read libraries.** Solid lines represent the number of poly(A) CS, and dashed lines represent the percentage of shared poly(A) CS over the overall poly(A) CS identified in each sequencing library.
